## Supplemental materials S3 for "A model of Hill-Robertson interference caused by purifying selection in a non-recombining genome"

**Supplementary File S1**

**1. The coalescent process with HRI**

Time is treated here as continuous. Assume that *j* alleles are present at the rescaled time *T* from the start of the process, with *T* = *t*/(*BN*_0_). Equations (9) of the main text imply that the rate of coalescence for this set of alleles is:

$\lambda_{j}\left( T \right)=\frac{j\left( j-1 \right)B}{2}$ $\exp(V_{a}Q_{T}^{2})$	(S1a)

where	$Q_{T}=\frac{V_{a}}{V_{m}}[1 -\exp(\frac{{N_{e}V}_{m}T}{V_{a}})]$	(S1b)

The expected time to the first coalescent event, at which the number of alleles decreases from *k* to *k* – 1, is given by the generalisation of Equation (12):

$\bar{T}_{k}=\int_{0}^{\infty} P_{nc, k}\left( T \right) dT$	(S2a)

where the probability of no coalescence by time *T* in the past is

$P_{nc,k}\left( T \right)\approx\exp[-\int_{0}^{T} \lambda_{k}\left( T \right) dX ]$	(S2b)

The p.d.f. for the time from the start of the process to the next coalescent event, when the number of alleles decreases to *k* – 2, is given by:

$\phi_{k-1}\left( T_{k-1} \right)=\int_{0}^{T_{k-1}} \psi_{k-1}\left( T_{k-1} | T_{k}=Y \right)\phi_{k}\left( Y \right)dY$	(S3a)

where the p.d.f. for *T_k_*_-1_ conditioned on *T_k_* = *Y* is:

$\psi_{k-1}\left( T_{k-1} | T_{k}=Y \right)=\lambda_{k-1}\left( T_{k-1} \right)\exp-\int_{X}^{T_{k-1}} \lambda_{k-1}\left( T \right)dT$	(S3b)

and the p.d.f for time *X* is given by:

$\phi_{k}\left( X \right)=\lambda_{k}\left( X \right)P_{nc,k}\left( X \right)$	(S3c)

The expected total time to the second coalescent event is thus:

$\bar{T}_{k-1}=\int_{0}^{\infty} T\phi_{k-1}(T) dT$	(S4)

This procedure can be repeated over successive coalescent events. Equation (S3a) generalises to:

$\phi_{k-j}\left( T_{k-j} \right)=\int_{0}^{T_{k-j}} \psi_{k-j}\left( T_{k-j} | T_{k+1-j}=X \right)\phi_{k+1-j}\left( X \right)dX$	(S5a)

and	$\bar{T}_{k-j}=\int_{0}^{\infty} T\phi_{k-j}(T) dT$	(S5b)

Let the rescaled time interval between *T_k – j_* and *T_k +_*_1_*_– j_* be Δ*T_k+_*_1_ *_– j_* (*j* = 2, …, *k* – 1), where $\bar{T}_{k}$ (*j* = 1) is given by Equations (S2).

The expected total length of the gene tree is given by:

$\bar{L}_{k}=k \bar{T}_{k}+\sum_{i=2}^{k-1} (k+1-i)\Delta\bar{T}_{k+1-i}$	(S6)

The probability of a segregating site in a sample of size *k* with a neutral mutation rate of *u* is given by:

$\bar{S}_{k}{=N}_{ec}u\bar{L}_{k}=BN_{0}u\bar{L}_{k}$	(S7a)

The expected value of Watterson’s *θ_w_* statistic is given by:

$\bar{\theta}_{w}=$ $\bar{S}_{k}/a_{k}$	(S7b)

where *a_k_* is the sum of the harmonic series (with terms 1/*i*) from *i* = 1 to *i* = *k* –1.

The ratio of the expected neutral nucleotide site diversity ($\bar{\pi}$) to $\bar{\theta}_{w}$ is thus given by:

$\frac{2B´N_{0}a_{k}}{BN_{0}\bar{L}_{k}}=\frac{2{B´a}_{k}}{B\bar{L}_{k}}$	(S8)

The corresponding value of the statistic Δ*θ_w_*, which provides a measure of the distortion of the site frequency spectrum (Becher et al. 2020), is thus:

$\Delta\theta_{w}=1-\frac{\bar{\pi}}{\bar{\theta}_{w}}=1-\frac{2B´a_{k}}{B\bar{L}_{k}}$	(S9)

An approximation to the expected value of Tajima’s *D* statistic is provided by:

$\hat{D}_{T}\approx\frac{L(\bar{\pi}-\bar{\theta}_{w})}{\sqrt{\bar{S}[\left( e_{1}-e_{2} \right)+e_{2}\bar{S}]}}$	(S10)

where $\bar{S}=$ $\bar{\theta}_{w}a_{k}$ is the mean number of segregating sites in a sample of *k* alleles, and the constants *e*_1_ and *e*_2_ are defined by Equations (36) and (37) of Tajima (1989).

The expected site frequency spectrum (SFS) of a sample of *k* alleles can be obtained using a simplified version of Equation (1.3) of Griffiths and Tavaré (1998) – an elementary derivation of this result is given in section 2 of this files. The probability that a derived mutation is present at a frequency of *i* in a sample of size *k* is given by:

$f\left( i \right)=\frac{\left( k-i-1 \right)!}{\left( k-1 \right)! \bar{L}_{k}} \sum_{j=2}^{j=k+1-i} \frac{\left( k-j \right)!k(k-1)}{\left( k+1-i-j \right)!}\Delta\bar{T}_{j}$	(S11)

The problem with implementing these formulae is that they involve multiple integrations over increasing numbers of variables as *k* increases, causing computational difficulties. Here, a sequence of stochastic integrations was used. For the first step (*k* to *k* – 1), in order to choose *T_k_* a random number *z* is chosen and compared to a set of numerical integrals representing the integrals *F_k_*(*Y*) of $\phi_{k}\left( X \right)$ in Equation (S3c) up to *X* = *Y*, using finely spaced values of *Y*. For *z* ≥ *F_k_*(*Y*), *T_k_* is set equal to *Y.* This value of *T_k_* is then used to determine the rates of coalescence over the next interval, and a set of numerical integrals *F_k_*_–1_ (*Y*) of $\phi_{k-1}\left( Y \right)$ determined from Equation (S3a). A new random z is chosen and the first value of *T_k_*_-1_ ≥ *z* chosen. This procedure is repeated until *T*_2_ has been set, requiring *k* – 1 random numbers in total to provide one set of values of the *T_k_.* The procedure is repeated *n* times, and the mean value and standard error of each *T_k_* determined.

**2. Derivation of the site frequency spectrum from the coalescent process**

Equation (1.3) of Griffiths and Tavaré (1998) for the neutral SFS in a general coalescent process uses classic results on occupancy problems from Chapter 2, §5 of Feller (1968) . Consider the problem of putting *r* balls into *n* cells. In the present notation, the *n* cells correspond to *j* ancestors at a given point in the past, and the *r* balls correspond to the *k* descendant alleles in the sample. The assumption is made that all arrangements are *a priori* equally probable; this is, of course, questionable when selection is acting.

Feller approached this problem by representing the cells by spaces between *n*+1 bars (I) and the balls by stars (*). For example, a configuration of 6 cells with occupancy numbers 3, 1, 0, 0, 0 and 4 is represented by I***I*I I I I****I. The genetic problem requires that each ancestor generates at least one descendant, so that no cell is empty. The *n* – 1 bars and *r* stars can appear in any order; it is assumed that these are all equally probable. There are thus *n* – 1 + *r* indistinguishable elements between the outer two bars, and the number of distinguishable arrangements is equivalent to the number of ways of placing *r* elements into these at random. But the condition that no cell is empty means that no two bars can be adjacent. There are *r* – 1 spaces between the *r* stars, of which *n* – 1 are to be occupied by bars, so that the number of possible arrangements with this restriction is $\left( \begin{aligned} r-1 \\ n-1 \end{aligned} \right)$, equivalent to $\left( \begin{aligned} k-1 \\ j-1 \end{aligned} \right)$ in my notation.

Now consider the case when *i* out of the *r* stars are in the same cell; this is equivalent to *i* descendant alleles coming from the same ancestral allele. There are then *r* – *i* stars to arrange into *n* – 1 cells, so that the above argument implies that there are $\left( \begin{aligned} r-i-1 \\ n-2 \end{aligned} \right)$ possible arrangements of this kind, equivalent to $\left( \begin{aligned} k-i-1 \\ j-2 \end{aligned} \right)$ in the genetic case. The probability of finding *i* descendants of the same ancestor from a generation when there are *j* ancestral alleles is thus:

$p_{k,j}\left( i \right)=\frac{\left( \begin{aligned} k-i-1 \\ j-2 \end{aligned} \right)}{\left( \begin{aligned} k-1 \\ j-1 \end{aligned} \right)}$	(S12)

This can be applied as follows to the probability of finding *i* derived mutations in a sample of *k* alleles under the infinite sites model, where only one mutation is segregating in the sample. Consider the time interval, Δ*T_j_* between *j* and *j* – 1 ancestral alleles in the gene tree. The probability of a mutation arising on one of the *j* alleles during this time interval is proportional to the product of *j* and the expectation of Δ*T_j_*, since there are *j* lines of descent present during this interval, so that the net probability of such a mutation is equal to the product of the mutation rate and ${j\Delta\bar{T}}_{j}$. The probability that this mutation generates *i* descendants in the sample is given by Equation (S12). The net probability of a mutation segregating in the sample is the product of the mutation rate and the total size of the tree, $\bar{L}_{k}$. Summing over all possible cases that can contribute to a mutation segregating among the *k* descendant alleles, the net probability of a mutation being present in *i* copies is given by:

$f\left( i \right)=\frac{1}{\bar{L}_{k}}\sum_{j=2}^{j=n+1-k} {jp}_{k,j}\left( i \right){\Delta\bar{T}}_{j}$	(S13)

Writing out the combinatorial expressions in Equation (S12) in terms of their component factorials, this expression simplifies to Equation (S11).

**3. Effects of *U* and *γ*_0_ on *B***

Some insights into the properties of HRI can be obtained by using Equation (7) to determine the partial derivative of *B* with respect to parameters of interest, notably *U*= *Lu* and *γ*_0_ = 2*N*_0_*s*. Equation (4) can be rewritten as:

$g\left( B,U, {\kappa,\gamma}_{0} \right)=ln\left( B \right)+ \frac{U}{s}{f(B,\kappa,\gamma_{0})}^{3}$	(S14a)

where

$f\left( B,\kappa, \gamma_{0} \right)= \frac{[1-\exp\left( -B\gamma_{0} \right)]}{[1+\kappa\exp\left( -B\gamma_{0} \right)]}$	(S14b)

For any parameter *λ*, implicit differentiation of Equation (S14a) gives:

$\partial_{\lambda}B=- \frac{\partial_{\lambda}g}{\partial_{B}g}$	(S15a)

and

$\partial_{\gamma}f=\frac{(1+\kappa)exp(\gamma)}{{[\kappa+\exp\left( \gamma\right)]}^{2}}$	(S15b)

We have:

$\partial_{B}g=B^{-1}+\frac{U}{s}\partial_{B}f^{3}=B^{-1}+\frac{3U}{s}\gamma_{0}f^{2}\partial_{\gamma}f$

$=B^{-1}+\frac{3\alpha_{0}(1+\kappa)\left[ \exp\left( \gamma\right)-1 \right]^{2}exp(\gamma)}{{[\kappa+exp(\gamma)]}^{4}}$	(S16)

where $\gamma={B\gamma}_{o}=2BN_{0}s, \alpha_{0}=2N_{0}U$,

It follows that $\partial_{B}g>0$ if *γ* ≠ 0, and so

$\partial_{U}B=-\frac{f^{3}}{s \partial_{B}g}$ < 0	(S17)

We thus expect *B* to be a decreasing function of *U*, the net mutation rate to deleterious alleles in the genomic region in question. This is, of course, what is expected intuitively and what is observed in the simulations and numerical results.

We next consider the partial derivative of *B* with respect to *N*_0_, keeping *s* constant:

$\partial_{N_{0}}g=\frac{U}{s}\partial_{\gamma}f^{3}={6UBf^{2} \partial}_{\gamma}f$	(S18)

From the above results, it follows that ${\partial B_{N}}_{0}$< 0, indicating that an enhanced efficacy of selection due to larger effective population size leads to a reduction in *B*.

The situation is more complex if *N*_0_*s* changes due to a change in *s* rather than *N*_0_, since its effect on the factor *U*/*s* as well as on *f* ^3^ needs to be considered:

$\partial_{s}g= -\frac{Uf^{3}}{s^{2}}+\frac{6U{BN}_{0}f^{2}}{s}\partial_{\gamma}f=-\frac{\alpha_{0}f^{2}}{s\gamma_{0}}(f-3\gamma\partial_{\gamma}f)$	(S19)

We thus have $\partial_{s}B>0$ if $f>3\gamma\partial_{\gamma}f$, i.e.,

$h =\left[ \kappa+\exp\left( \gamma\right) \right]\left[ 1-\exp\left( -\gamma\right) \right]-3\left( 1+\kappa\right)\gamma>0$	(S20)

When γ >> 1, this condition is satisfied, corresponding to the result that *B* is a decreasing function of *U*/*s* in the classical background selection case described by Equation (6). When *γ* is sufficiently small, it is violated, and *B* increases as *s* decreases, with *B* = 1 when *s* = 0. The partial derivative of *h* with respect to *γ* is:

$\partial_{\gamma}h= \kappa\exp\left( -\gamma\right)+\exp\left( \gamma\right)-3(1+\kappa)$	(S21)

A sufficient condition for $\partial_{\gamma}h>0$ is $\gamma>ln[3\left( 1+\kappa\right)]$; this can be met for quite modest value of *γ*. For example, with *κ* = 3, this condition is satisfied when *γ* > 2.485. After this point, *h* will always increase with *s* and eventually becomes positive.

These results suggest that *B* is a decreasing function of *s* for small *N*_0_*s*, if is held constant and *s* is varied; it eventually becomes an increasing function of *s* when *N*_0_*s* is is sufficiently large.

**4.** **Comparisons with the simulation results of Comeron and Kreitman (2002)**

The model of Comeron and Kreitman (2002) (CK) assumes a non-recombining chromosome consisting of equal sized blocks of selected and neutral sites. This is a diploid model, so 2*N* in their model corresponds to *N* in a haploid model. They also parameterize selection by a selection coefficient *s* against homozygotes. Their *α* = *Ns* thus corresponds to *Ns*/2 in our haploid model. In our notation, the mutation rate from preferred to unpreferred states is *u* and the reverse mutation rate is *v*, with *u* = *κv*. In their notation, these rates are *w* and *v*, respectively, and their mutational bias parameter is *γ* = *v*/(*u* + *v*). The relation between *κ* and *γ* can be found by using *γ* (*u* + *v*) = *v*, implying that *v* = *γu*/(1 – *γ*); *u* = (1 – *γ*)*v*/*γ*. This implies that *κ* = (1 – *γ*)/*γ*. For *γ* = 0.45, as used by CK, *κ* = 1.222. In our notation, their scaled mutation parameter *β* = *N*(*u* + *v*)/2 = *Nu*[1 + 1/*κ*]/2 = *Nu*[1 + *κ*]/(2*κ*). Thus, we have *Nu* = 2*κβ*/(1 + *κ*).

| Genome size | Simulated ratio of selected to neutral *π* | Theoretical ratio of selected to neutral *π* | Simulated  *B* for neutral *π*  (*B*´) | Theoretical  *B* for neutral *π*  (*B*´) | BGS value  of *B* |
| --- | --- | --- | --- | --- | --- |
| *s* = 2.5 x 10^-4^ |  |  |  |  |  |
| 125 | 0.96 | 0.897 | 0.965 | 0.942 | 0.0639 |
| 250 | 0.80 | 0.859 | 0.84 | 0.908 | 0 |
| 500 | 0.75 | 0.814 | 0.75 | 0.864 | 0 |
| 2500 | 0.70 | 0.719 | 0.71 | 0.730 | 0 |
| *s* = 5.0 x 10^-4^ |  |  |  |  |  |
| 125 | 0.86 | 0.766 | 0.91 | 0.876 | 0.253 |
| 250 | 0.725 | 0.735 | 0.76 | 0.829 | 0.064 |
| 500 | 0.675 | 0.698 | 0.725 | 0.758 | 0.004 |
| 2500 | 0.3 | 0.611 | 0.64 | 0.601 | 0 |
| *s* = 1.0 x 10^-3^ |  |  |  |  |  |
| 125 | 0.70 | 0.589 | 0.875 | 0.802 | 0.503 |
| 250 | 0.61 | 0.593 | 0.68 | 0.722 | 0.253 |
| 500 | 0.55 | 0.583 | 0.625 | 0.642 | 0.064 |
| 2500 | 0.475 | 0.535 | 0.50 | 0.477 | 0 |
| *s* = 2.5 x 10^-3^ |  |  |  |  |  |
| 125 | 0.425 | 0.271 | 0.80 | 0.820 | 0.760 |
| 250 | 0.37 | 0.321 | 0.63 | 0.689 | 0.577 |
| 500 | 0.35 | 0.393 | 0.52 | 0.539 | 0.333 |
| 2500 | 0.32 | 0.447 | 0.38 | 0.339 | 0.004 |

**Table S1 Comparisons of CK simulation results with theoretical predictions**

Simulation results taken from Fig. 1 of Comeron and Kreitman (2002). Diploid N = 500.

Note that the neutral *π* in the absence of HRI is 0.0198 with their parameters.

The CK results for the second column are interpreted as the ratios of the diversities at selected sites to those at neutral sites.

No correction for the effect of LD on *π* at selected sites is made here.

**5. Comparisons with the simulation results of McVean and Charlesworth (2000)**

The simulations of McVean and Charlesworth (2000) (MC) used a diploid model with variable rates of recombination, with varying numbers of sites under selection, no mutational bias, and no neutral sites. Only the zero recombination rate results are considered here. With their parameterization of selection, their 4*N_e_s* in the absence of HRI is equivalent to our 4*N*_0_*s*, and their scaled mutation rate of 4*N_e_u* to our 2*N*_0_*u*. They parameterized the effect of selection on the level of adaptation by using a “codon bias” index, which is equivalent to $c=2\bar{p}-1$.

$2\bar{p}-1=2\left( 1+\kappa exp-\gamma\right)^{-1}-1=(1-\kappa exp-\gamma){(1+\kappa exp-\gamma)}^{-1}$	(S22)

Comparison with the expression for *π* for selected sites (Equation 2a), shows that, provided that the same value of *B* applies to both diversity and $\bar{p}$, the ratio of both *c* and *π* for cases with and without HRI are given by:

$\frac{(1-\kappa exp-\gamma)(1+\kappa exp-\gamma_{0})}{(1+\kappa exp-\gamma)(1-\kappa exp-\gamma_{0})}$	(S23)

Table S2 shows comparisons of the simulation results for the ratios of *c* and *π* with HRI to the predicted values of the ratios. Table S3 shows simulated values of *B*, obtained using Equation (6), with the corresponding predictions.

**Table S2 Comparison of MC simulation results on “codon bias” and diversity at selected sites with theoretical predictions**

| Genome sizes | Simulated ratio of codon bias  to single-locus value | Theoretical ratio of codon bias to  single-locus value (also applies to  ratio of *π* at selected sites) | Simulated ratio of selected site *π* to single-locus value | Theoretical  ratio of selected site *π* using  neutral *B*  (*B*´) | *B* for neutral  Diversity  (*B*´) | BGS value  of *B* |
| --- | --- | --- | --- | --- | --- | --- |
| *γ* = 1 |  |  |  |  |  |  |
| 500 | 0.75 | 0.634 | 0.91 | 0.822 | 0.800 | 0 |
| 1000 | 0.675 | 0.550 | 0.83 | 0.766 | 0.740 | 0 |
| 5000 | 0.51 | 0.376 | 0.725 | 0.620 | 0.590 | 0 |
| *γ* = 2 |  |  |  |  |  |  |
| 500 | 0.675 | 0.562 | 0.86 | 0.781 | 0.685 | 0 |
| 1000 | 0.60 | 0.478 | 0.84 | 0.722 | 0.618 | 0 |
| 5000 | 0.425 | 0.317 | 0.63 | 0.572 | 0.467 | 0 |
| *γ* = 4 |  |  |  |  |  |  |
| 500 | 0.70 | 0.620 | 0.92 | 0.840 | 0.565 | 0.007 |
| 1000 | 0.625 | 0.523 | 0.93 | 0.786 | 0.496 | 0 |
| 5000 | 0.45 | 0.340 | 0.70 | 0.636 | 0.357 | 0 |

Simulation results are from Figure 3 of McVean and Charlesworth (2000).

**Table S3 Comparison of MC simulation results on B values for selected sites with theoretical predictions**

| γ | *B* for N_e_s: simulated | *B* for N_e_s: theoretical |
| --- | --- | --- |
| 1 | 0.59 ± 0.05 | 0.627 |
| 2 | 0.46 ± 0.03 | 0.381 |
| 4 | 0.34 ± 0.02 | 0.277 |
| 10 | 0.24 ± 0.01 | 0.192 |
| 20 | 0.20 ± 0.01 | 0.182 |

Simulation results are from Table 1 of McVean and Charlesworth (2000).

**6. Distributions of the summary statistics obtained from the simulations**

**Figure S1** The distributions of the summary statistics $\bar{p}$ , *B*, *B*´ and $\Delta\theta_{w}^{'}$, obtained from the simulations described in the section *Results: Levels of adaptation and neutral diversity under HRI, Fits to new simulation results.*

Each box represents one set of 200 replicate simulations. The fat horizontal lines indicate the medians. The top and bottom lines of each box indicate the 75% and 25% quantiles, between which lies the 'interquartile range'. The dashed 'whisker' lines indicate the total spread of the data, except that each is limited to be no longer than 1.5 times the interquartile range. Data points further away are shown as circles.

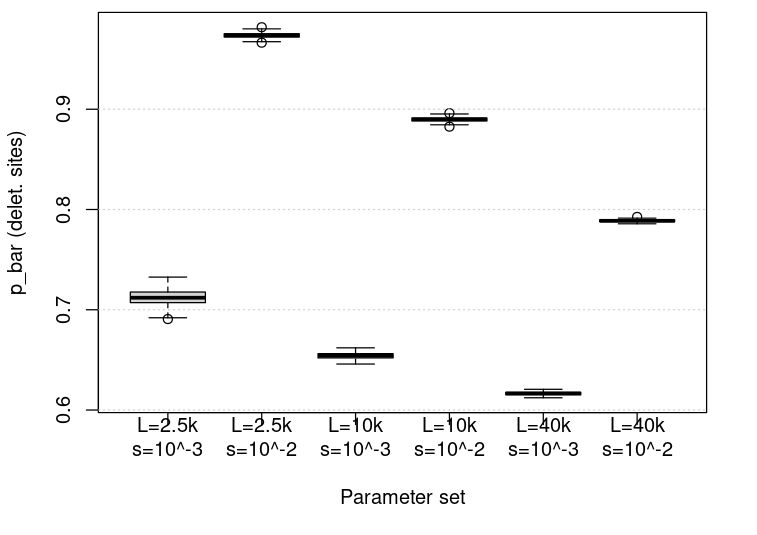

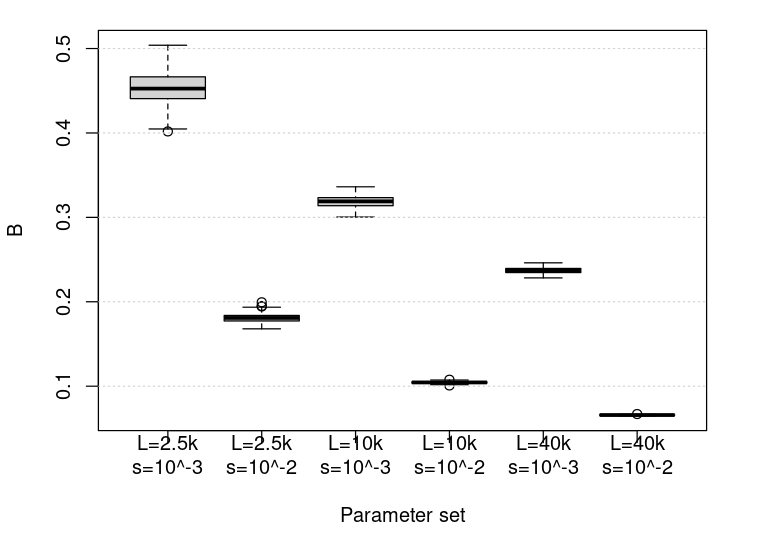

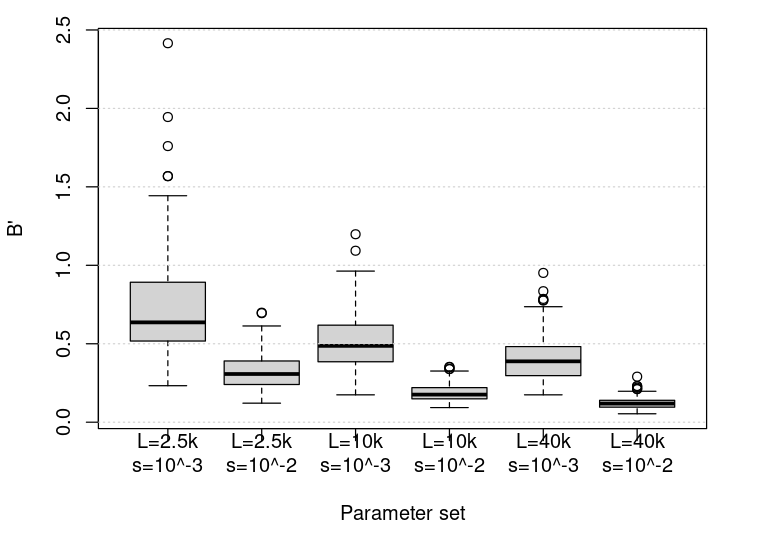

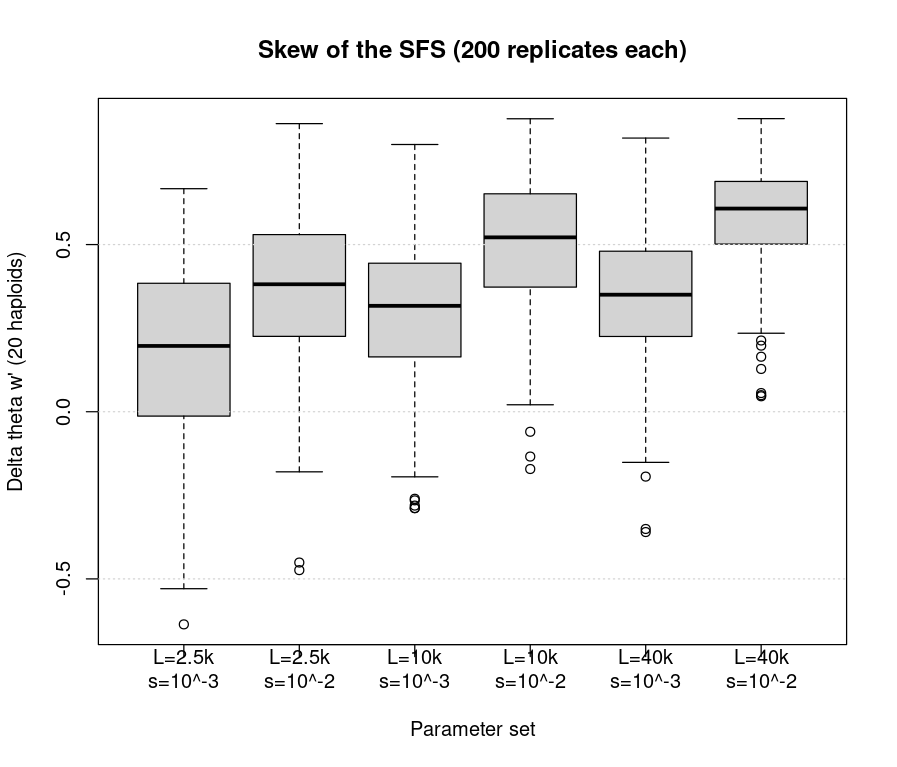

**7. Comparisons of different site frequency spectrum summary statistics from the simulation results**

**Table S4. Mean values from the simulations of (upper entries in each cell) and Tajima’s**

***D* (lower entries) for neutral sites, showing the effects of different sample size**

| Sample  size | *L* = 2500  *γ* = 2 | *L* = 2500  *g* = 20 | *L* = 10000  *g* = 2 | *L* = 10000  *g* = 20 | *L* = 40000  *g* = 2 | *L* = 40000  *g* = 20 |
| --- | --- | --- | --- | --- | --- | --- |
| 10 | 0.148  (0.099,0.196)  –0.317  (–0.422,  –0.213) | 0.321  (0.278,0.365)  – 0.681  (–0.772,–0.589) | 0.291  (0.217,0.303)  – 0.565  (–0.658,  –0.471) | 0.469  (0.436,0.502)  – 1.012  (–1.08,  –0.940) | 0.295  (0.255,0.335)  –0.642  (–0.730,–  0.555) | 0.532  (0.500,0.563)  –1.155  (–1.224,–1.087) |
| 20 | 0.173  (0.135,0.211)  – 0.461  (–0.562,–  0.360) | 0.391  (0.361,0.421)  – 1.026  (–1.105,–0.947) | 0.290  (0.257,0.323)  – 0.783  (–0.871,–0.693) | 0.510  (0.4483,0.538)  – 1.366  (–1.44,–1.29) | 0.357  (0.328,0.386)  –0.965  (–1.04,–0.886) | 0.573  (0.552,0.594)  –1.546  (–1.602,–1.490 |
| 40 | 0.186  (0.151,0.221)  – 0.542  (–0.644,  –0.440) | 0.423  (0.397,0.449)  – 1.218  (–1.292,–1.144) | 0.307  (0.278,0.335)  – 0.905  (–0.989,–0.821) | 0.556  (0.539,0.581)  – 1.643  (–1.70,–1.58) | 0.377  (0.353,0.402)  –1.117  (–1.190,–1.045) | 0.629  (0.613,0.646)  –1.881  (–1.910,–1.811) |
| 80 | 0.184  (0.148,0.221)  – 0.550  (–0.656,  –0.445) | 0.454  (0.429,0.479)  –1.349  (–1.422,–1.275) | 0.333  (0.306,0.360)  – 1.012  (–1.09,–0.931 | 0.605  (0.588,0.623)  – 1.834  (–1.887,–1.781) | 0.400  (0.377,0.423)  –1.222  (–1.293,–1.151) | 0.671  (0.657,0.685)  –2.045  (–2.088,–2.002) |

95% confidence intervals of the means are shown in brackets.

**8. Increase in the probability of high frequency derived variants at the extreme end of the site frequency spectrum**

|  | Sample of 20 |  | Sample of 80 |  |
| --- | --- | --- | --- | --- |
| Parameters | Simulations | Polanski et al. | Simulations | Polanski et al. |
| *L* =2500, *γ*_0_ = 2 | 1.24 | 0.93 | 1.93 | 0.98 |
| *L* =2500, *γ*_0_ = 20 | 1.18 | 0.94 | 1.53 | 0.99 |
| *L* =10000, *γ*_0_ = 2 | 1.29 | 0.93 | 1.45 | 0.98 |
| *L* =10000, *γ*_0_ = 20 | 1.54 | 0.94 | 1.21 | 0.99 |
| *L* =40000, *γ*_0_ = 2 | 1.34 | 0.93 | 2.70 | 0.98 |
| *L* =40000, *γ*_0_ = 20 | 1.20 | 0.94 | 0.85 | 0.99 |

The entries in each cell are the ratios of the frequencies of 19 to 18 derived variants (sample size 20) and 79 versus 78 derived variants (sample size 80) from the simulations or from the theoretical predictions based on Polanski et al. (2003)
